# Comprehensive analysis of gene regulatory dynamics, fitness landscape, and population evolution during sexual reproduction

**DOI:** 10.1101/2022.05.11.491509

**Authors:** Kenji Okubo, Kunihiko Kaneko

**Affiliations:** Research Center for Integrative Evolutionary Science, The Graduate University for Advanced Studies; Center for Complex Systems Biology, Universal Biology Institute, University of Tokyo; Niels Nohr Institute, University of Copenhagen

**Keywords:** fitness landscape, gene regulatory network, sexual reproduction

## Abstract

The fitness landscape is a critical concept in evolutionary biology and genetics that depicts fitness in the genotype space and visualizes the relationship between genotype and fitness. However, the fitness landscape is challenging to characterize because the quantitative relationships between genotype and phenotype and their association to fitness has not been comprehensively well described. To address this challenge, we adopted gene regulatory networks to determine gene expression dynamics. We analyzed how phenotype and fitness are shaped by the genotype in two-gene networks. A two-by-two matrix provided the two-gene regulatory network in which a vector with two angle values (Θ) was introduced to characterize the genotype. Mapping from this angle vector to phenotypes allowed for the classification of steady-state expression patterns of genes into seven types. We then studied all possible fitness functions given by the Boolean output from the on/off expression of the two genes. The possible fitness landscapes were obtained as a function of the genetic parameters Θ. Finally, the evolution of the population distribution under sexual reproduction was investigated in the obtained landscape. We found that the distribution was restricted to a convex region within the landscape, resulting in the branching of population distribution, including the speciation process.

## Introduction

The fitness landscape in evolutionary biology describes fitness as the height of the genotype space and intuitively visualizes the relationship between the genotype and fitness. The fitness landscape provides a significantly simplified picture of evolutionary biology and genetics and is relevant to study evolvability, evolutionary pathways, the effects of multiple mutations, and speciation.

However, it is difficult to obtain suitable fitness landscapes. In a fitness landscape, the genotype is represented by a set of parameters that can be mapped onto the fitness. According to the landscape, how fitness changes depends on the genetic parameters that are prescribed. However, there are two fundamental problems with this approach. It is necessary to determine the genetic parameters that describe landscape axes. Genetic information can be written as the DNA sequence and how continuous parameters are derived from this genetic sequence is not trivial because a slight change in the sequence could significantly change the phenotype. Therefore, the derivation of continuous and measurable genetic parameters that create the fitness landscape must be addressed. Second, fitness is a function of phenotype and is not computed from the genotype directly. Usually, the expression of RNAs and proteins is a complex and dynamic process that determines a resultant phenotype. In other words, the relationship between genotype and phenotype depends on complex gene expression dynamics. Fitness, then, is a function of dynamically regulated phenotype.

We adopted gene regulatory networks (GRN) that describe gene expression dynamics to address these questions about the fitness landscape. First, we showed that the degree of activation or inhibition in expression dynamics continuously defines genotype parameters. Second, phenotypes, that is, protein expression levels, are determined by gene expression dynamics; whereas, genomes provide the gene regulatory network. Thus, a complex genotype–phenotype relationship was obtained. Then, fitness was defined by the phenotypes, and thus, represented as a function of the introduced genetic parameters. In fact, the evolution of GRN has been studied(Glass and Kauffman (1973); Mjolsness *et al*. (1991); Salazar-Ciudad *et al*. (2001, 2000); Kaneko (2006)) extensively with respect to the robustness or the phenotypic plasticity(Martin and Wagner (2009); Wagner (2013); Azevedo *et al*. (2006); Glass and Kauffman (1973); Mjolsness *et al*. (1991); Salazar-Ciudad *et al*. (2001, 2000); Kaneko (2006); Okubo and Kaneko (2021a,b); Kaneko (2007); Swain *et al*. (2002); Ou *et al*. (2008); Furusawa *et al*. (2005); Ayroles *et al*. (2015); Cubillos *et al*. (2014); Chapal *et al*. (2019); Miller *et al*. (2015); Kaneko and Kikuchi (2020); Nagata and Kikuchi (2020)).

While there are extensive studies of evolution in the fitness landscape(Soyer and Bonhoeffer (2006); Neyfakh *et al*. (2006); Ho and Zhang (2016); Orlenko *et al*. (2016); Yubero *et al*. (2017); Friedlander *et al*. (2017); Cuypers *et al*. (2017); Orlenko *et al*. (2017); Schiffman and Ralph (2022); Hether and Hohenlohe (2014)), the relationship between the global fitness landscape and GRN 2 Comprehensive analysis of gene networks structures remains uncharacterized. In particular, Heather and Hohenlohe (Hether and Hohenlohe (2014)) classified GRN dynamics into six cases. However, the global change in gene expression dynamics due to GRN changes and the classification of the fitness landscape need to be further investigated by comprehensively considering all classes of possible GRNs and their relationship to phenotypes and fitness.

Generally, GRNs and their dynamic interactions with many regulatory genes are too complex to analyze. Here, we consider a GRN with two genes, which provides a straightforward and basic system to study comprehensively the dynamics that create a genotype–phenotype relationship. Here, two-node GRNs are represented by 2×2 matrices. By using on/off expression dynamics, we could evaluate all the possible GRN structures by introducing two-dimensional parameters specifying the matrix.

Still, there can be many types of fitness functions for given expressions; in most theoretical studies, however, a specific fitness function is selected depending on the purpose of the study. Here, we studied all possible fitness functions that depended on the expression of the two genes (i.e., Boolean functions). The possible fitness functions were limited to 16 types, all of which were investigated to obtain the possible fitness landscape. Once the fitness landscape was obtained, the genome distribution of the GRN parameter was computed. The distribution was concentrated on a higher-fitness region; however, the robustness against recombination or sexual reproduction (Martin and Wagner (2009); Azevedo *et al*. (2006); Kim and Fernandes (2009); Okubo and Kaneko (2021a); Omholt *et al*. (2000)) will impose further restrictions on the genome distribution, as offspring from the parent selected from the high-fitness region could fall into a non-fitted region. Hence, it was crucial to determine how stability against recombination shaped the genome distribution depending on the fitness landscape.

This paper consists of two parts. The first part analyzes the dynamics of the two-gene network and the second part discusses the convex regionalization of the population by sexual reproduction based on the fitness landscape. (See Fig.1 for the flow used in this study).

**Figure 1.**
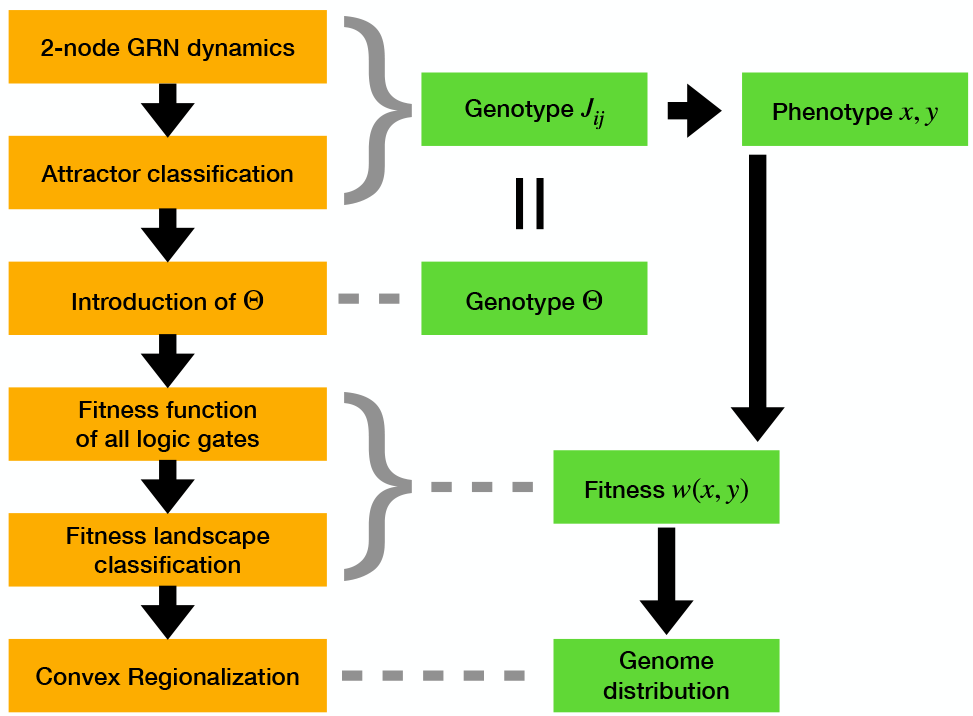
A flow chart in this paper to obtain the genotype– phenotype relationship and fitness landscape.

In the first part, the phenotype and fitness are obtained from the genotype in a two-gene network. Because the two-gene network has only a few possible steady states, we classified these steady states by introducing the angle vector Θ = (*θ*_*x*_,*θ*_*y*_) from the 2×2 gene regulation matrix. Θ provides the characteristic parameters of the genotype. Next, we considered fitness as a function of the on/off expression patterns to describe the fitness landscape as a function of Θ.

The second part discusses the evolution of population distribution by sexual reproduction within all possible fitness landscapes. In particular, when the high-fitness region consisted of two disjointed parts, we found that the speciation of the two groups occurred by sexual recombination, whereas convex regionalization from the non-concave fitted region was also demonstrated. Finally, we discuss the relationship between the fitness landscape and the convex regionalization of epistasis, sexual reproduction, and speciation.

### Dynamics of the two-gene regulatory network and fitness landscape

#### Two-gene regulatory network model

Our two-gene regulatory networks model assumed that there were two genes *X* and *Y*. Let *x* and *y* be the concentrations of proteins transcribed and translated from each gene, respectively Fig.2. Proteins X and Y are produced from the transcription of genes *X* and *Y*, respectively, and bind to the promoter regions of *X* and *Y* (represented by the squares on the line in Fig.2. When proteins X and Y bind to a gene promoter, they either promote or inhibit transcription of that gene. When protein *X* regulates the transcription of gene *X*, the degree of regulation is represented by *J*_*xx*_. Similarly, the regulation of gene *X* by *Y* is represented by *J*_*xy*_, regulation of gene *Y* by *X* by *J*_*yx*_, and regulation of gene *Y* by *Y* by *J*_*yy*_. These are collectively denoted by *J*_*ij*_. The value of *J*_*ij*_ can take any real value; thus, when *J*_*ij*_ is positive, transcription is promoted, and when *J*_*ij*_ is negative, it is inhibited.

**Figure 2.**
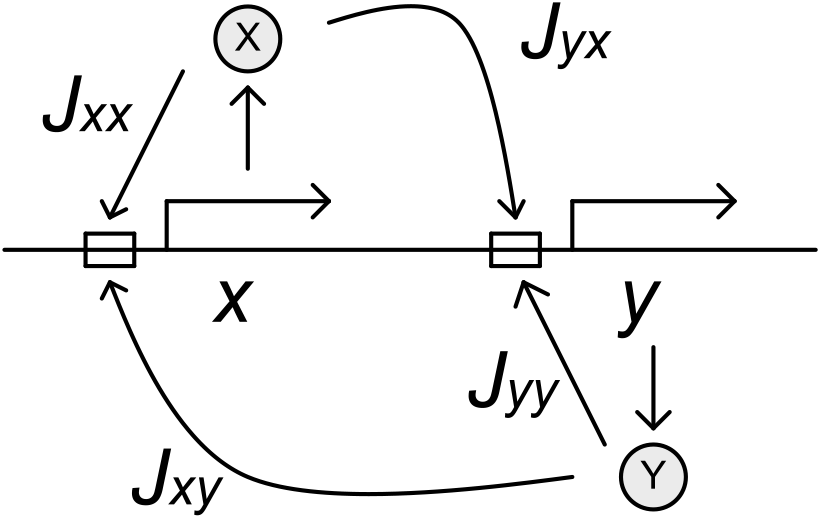
Regulatory network of two genes. The arrows represent the transcribed regions and the boxes represent the promoter regions. The mRNA is transcribed from each of the X- and Y-transcribed regions, from which proteins X and Y are synthesized. These proteins bind to the promoter regions of the two genes and regulate their transcription. The translation is assumed to occur very quickly and is therefore, omitted from this figure. Because proteins X and Y regulate genes *x* and *y*, there are four interactions. The magnitudes of these interactions are indicated by *J*_*xx*_, *J*_*xy*_, *J*_*yx*_, and *J*_*yy*_

When a protein binds to a promoter, gene transcription is promoted or inhibited. For simplicity, we assumed that a gene is transcribed when the sum of the effects of molecular regulation exceeds a threshold value. The activation function was given by the sigmoid function 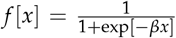. The degree by which gene *m* expression is regulated by protein N transcribed by geneare reported by *n* as *J*_*mn*_, with the dynamics of the expression level and protein concentration of X and Y represented by

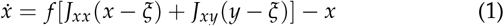

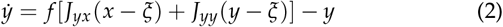

−*x* and −*y* on the right-hand side of the second term represent protein X and Y degradation, respectively. Here, the expression threshold was set at 0.5, to make the expression and nonexpression states symmetric for later fitness function simplicity. *β* of the sigmoid function was set to 100 to make the Hill function 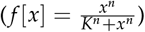 sharp.

#### Attractors of gene-expression dynamics

Considering the relationship between genotype and phenotype, we focused on where the expression levels *x* and *y* converged to a fixed point or cycle. We investigated how phenotype (*x, y*) was determined depending on the genotype, *J*_*ij*_(*i, j* = *x, y*).

To investigate the fixed point, we first obtained the nullclines of *x* and *y*, respectively, which were curves that satisfied 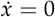 = 0 or 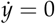 = 0 in Eq.(2). The nullclines were represented by the equation:

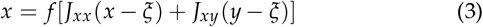

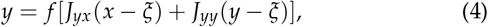

which are illustrated in Fig.3. Since *β* >> 1, *f* [*x*] approaches a step function, so that *x* = *f* [*x*] is satisfied either at *x* ≈ 1 or *x* ≈ 0, as well as *x* ≈ *ζ* = 0.5. In this paper, for simplicity, these *x* ≈ 1 and *x* ≈ 0 states are written as just *x* = 1 and *x* = 0 considering *β* >> 1.

**Figure 3.**
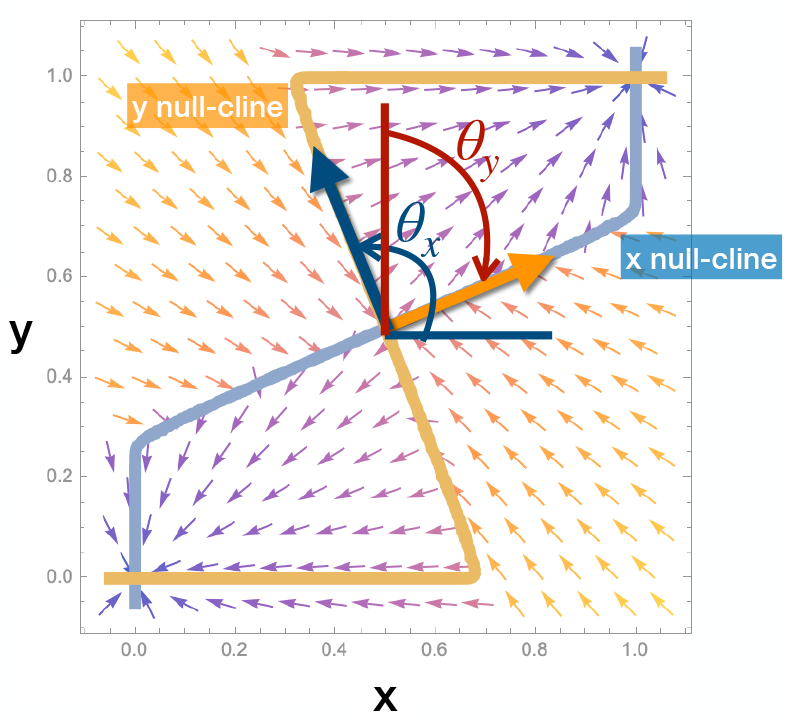
Nullcline normal vector. First, the nullclines were obtained from 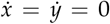 in Eq.(1) and Eq.(2). The blue line represents the nullcline for x and orange that for y. The intersection of these two null lines was a fixed point. The shape of the nullcline was characterized by normal vectors that indicate the directions of the nullclines around (0.5,0.5) and are denoted as (*J*_*xx*_, *J*_*xy*_) or (*J*_*yx*_, *J*_*yy*_). Furthermore, the vector field around the fixed point assisted in determining whether the fixed point was a stable fixed point (i.e., when the vector field around the fixed point was directed toward the fixed point from any direction) or an unstable fixed point (i.e., when the vector field around the fixed point is directed away from it). In this figure, (0,0), (0.5,0.5), and (1,1) are nullcline intersections that are fixed points, whereas only (0,0) and (1,1) are stable fixed points and (0.5,0.5) is an unstable fixed point.

#### Classification of two-gene expression dynamics by attractors

For given gene regulatory networks, phenotypes determined by attractors in gene expression dynamics can be organized into seven classes (Fig.4). The attractor types, their numbers, and their configurations are classified based on the value of *J*_*ij*_ as follows:

1. Equal expression level of *x, y* (S, symmetric or synchronized) *x* = 1, *y* = 1, or *x* = 0, *y* = 0 is the fixed-point attractor depending on the initial conditions. We referred to this as dynamics-type-S.
2. Different expression level of *x, y* (A, antagonistic) *x* = 1, *y* = 0 or *x* = 1, *y* = 0 is the fixed-point attractor depending on the initial conditions. We referred to this dynamic as type-D.
3. Same or different expression level of *x, y* (Q, quad) All possible four cases with *x* = 0 or 1, *y* = 0 or 1 give the fixed-point attractor depending on the initial conditions and were referred to as dynamics-type-Q.
4. Intermediate expression level of *x* (Cx, continuous for *x*) *x* = *α, y* = 0 or *x* = *α, y* = 1 is the fixed-point attractor, where 0 *< α <* 1, depending on the initial conditions and were defined as dynamics-type-Cx.
5. Intermediate expression level of *x* (Cy, intermediate for *y*) *x* = 0, *y* = *α* or *x* = 1, *y* = *α* is the fixed-point attractor, where 0 *< α <* 1, depending on the initial conditions. We defined this a dynamics-type-Cy.
6. Half expression level for *x, y* (H, half) *x* = 0.5 and *y* = 0.5 is the fixed-point attractors that were observed all initial conditions were defined as dynamics-type-H.
7. Periodic expression *x, y* (P, periodic) When the limit cycle was obtained in all initial conditions, we referred to this as dynamics-type-P.

**Figure 4.**
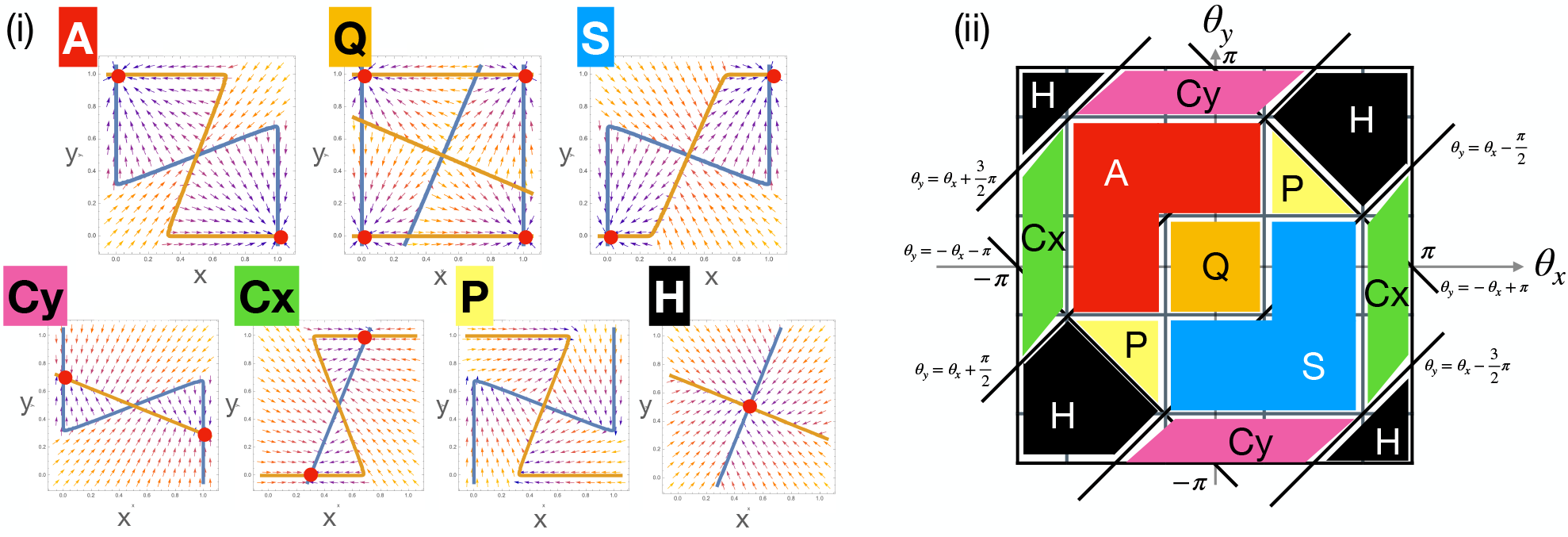
Fixed point classification of the Θ torus. (i)Each type of dynamics and nullclines in the two-gene regulatory network is represented. The Θ values for each type in the figure are 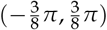 for (A), 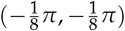 for (Q), 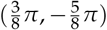 for (S), 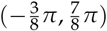 for (Cy), 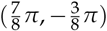 for (Cx), 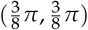 for (P), and 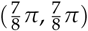 for (H). (ii) Classification of each dynamic category was based on (*θ*_*x*_, *θ*_*y*_). The boundaries between the categories in the figure are 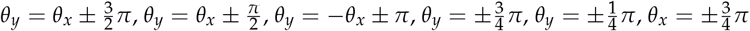, and 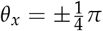;, respectively.

#### Introduction of Θ

We were able to classify steady states that corresponded to phenotypes and introduced a vector that characterized the genotype. Originally, the genotype in the model was represented by a *J*_*ij*_ in the 2×2 real matrix. However, the GRN that adopted a sigmoid function only uses two states, 0, 1, beside 0.5, which reduced the dimension of the genotype. This two-dimensional parameter represented the genotype’s declination angle, Θ.

We then focused on the shape of the nullcline in Fig.3. The shape was determined by the direction of the line segment that passed through point (0.5,0.5), which determined the attractor. Therefore, the change in the dynamics were due to the change in *J*_*ij*_ and could be specified in the direction of normal vectors of the *J*_*ij*_ row vectors. When examining the direction of this line segment, one of the normal vectors of this line segment (Fig.3) pointed in the same direction as the row vector of *J*_*ij*_, that is, (*J*_*xx*_, *J*_*xy*_) or (*J*_*yx*_, *J*_*yy*_). Here, only the direction of the row vectors mattered because the magnitude of the vectors was related to the shape of the nullcline. Hence, the shape of the nullcline was described by the angles of the *J*_*ij*_ (Θ) row vectors. In other words, as the angle between a line extended from the origin and the row vector, (*J*_*xx*_, *J*_*xy*_) or (*J*_*yx*_, *J*_*yy*_), we could take advantage of the symmetry between *x* and *y* to define the counterclockwise angle between the positive part of the x-axis and (*J*_*xx*_, *J*_*xy*_) as *θ*_*x*_, and the counterclockwise angle between the positive part of the y-axis and (*J*_*yx*_, *J*_*yy*_) as *θ*_*y*_. They were defined by

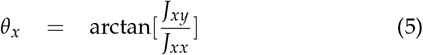

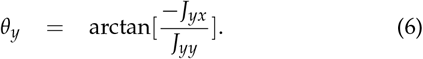

The dynamics and attractors were determined by using Θ = (*θ*_*x*_,*θ*_*y*_)

#### Θ torus and attractor classification

Using *θ*_*x*_ and *θ*_*y*_, we could explicitly classify the attractors of the teo-gene regulatory dynamics (Fig.4). This section introduces the *θ* torus, which had a finite range of *θ*_*x*_ and *θ*_*y*_ and specified each attractor type.

First, the behavior of the gene expression dynamics was determined by the direction of the *J*_*ij*_, Θ row vector. Here, the phenotype was visualized by introducing the Θ torus with *θ*_*x*_ on the horizontal axis and *θ*_*y*_ on the vertical axis, where *θ*_*x*_ *θ*_*y*_ were cyclic with mod 2*π*, as in Fig.4. Therefore, Θ torus represented the attractor types. Each of the types (S, D, Q, Cy, Cx, H, and P) was characterized by the nullclines shown in Fig.4A. Note that Θ determined the directions in the nullcline near point (0.5,0.5). The attractor determined by the two nullclines could change depending on the relative positions of *θ*_*x*_ and *θ*_*y*_. Then, S, D, Q, Cy, Cx, H, and P were classified depending on (*θ*_*x*_, *θ*_*y*_) as presented in Fig.4B. For the derivation of the boundaries for each classification, see Supplementary Material S1. This diagram shows the relationship from Θ to the attractors in (*x, y*) and the one between genotype and phenotype.

#### Defining the fitness function

We obtained the fitness landscape for Θ torus by first defining the fitness. Here, the phenotype was determined by the stationary expression levels of *x* and *y*. Here, we consider the case that the fitness takas maximal either at each expression level 0 or 1 (i.e., the fitness depends monotonically on the combination of each expression level, so that the fitness takes maximum at 0 or 1). Hence we focused on the states with values of 0 or 1; therefore, the fitness was described by a combination of the four-state input of (*x, y*) = (0, 0), (0, 1), (1, 0), (1, 1) and their two-state output. We assumed that binary fitness (fitted or non-fitted) was based on the binary phenotype. Hence, the fitness function was described by a Boolean (logical) function with two binary inputs and one binary output. In this case, 2^4^ = 16 possible fitness degree functions existed, which were reduced to five fitness functions using the symmetry of the model and identifying logical operations. (See Supplementary Material S2).

As for the fitness function *w*(*x, y*), we introduced a continuous function to satisfy the Boolean function such that *w*(*x, y*) for *x, y* = (0, 1) was either 0 or 1, and determined that the simplest form for *w*(*x, y*) had intermediate values between 0 and 1. We characterized four typical fitness functions here. Other cases including NEUTRAL are described in Supplementary Material S3.

##### AND: requires expression of both genes

This function used a maximum value of 1 when both *x* and *y* were set to 1.

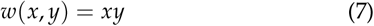

This corresponded to the case where *x* and *y* expression was required for survival (e.g., the formation of complexes by the proteins X and Y).

##### X ONLY: requires only X, whereas Y expression is neutral

In this function, fitness referred only to *x* and thus, if *x* was 1, fitness returned a maximum value of 1.

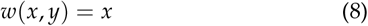

This corresponded to the selection pressure affecting only the expression level of *x* (e.g., protein X is functionally dominant or protein Y had a neutral function). When the fitness depended only on the expression of *y*, it presented similarly to that of *w*(*x, y*) = *y*.

##### XOR: requires the expression of only X or Y

When (*x, y*) was (1, 0) or (0, 1), the fitness had a maximum value of 1. By contrast, when (*x, y*) was (0, 0) or (1, 1), the fitness had a minimum value of 0. As the simplest form,

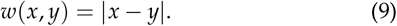

This function required that the expression of only one gene, *x* or *y*, was necessary for survival, but fitness was lost if both genes were expressed (e.g., switching the two pathways by the inputs).

##### OR: requires the expression of either one of the two genes

If at least one of *x, y* is 1, then the fitness had a maximum value of 1.

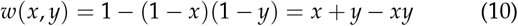

This corresponded to the case in which the expression of *x* or *y* was needed for survival (e.g., both X and Y proteins have similar functions).

#### Classification of the fitness landscape

We next explored the fitness landscape of genotype spaces that depended on the fitness function represented by Θ torus. Here, for a given Θ, we obtained the attractor of (*x, y*) for a fixed initial expression (*x*_0_, *y*_0_) = (0.05, 0.5) ((*x*_0_, *y*_0_) = (0.18, 0.18) for the XOR. Depending on the fitness function, we defined (AND, X ONLY, XOR, *etc*.) to obtain the fitness landscape for Θ, as shown in the diagram presented in Fig.4B, which contains information about the complex genotype–phenotype relationship.

We defined four typical landscapes (ONE RECTANGLE, TWO RECTANGLES, L-SHAPE, and ONE BAND in Fig.5), as well as additional landscapes that are described in Supplementary Material S3.

**Figure 5.**
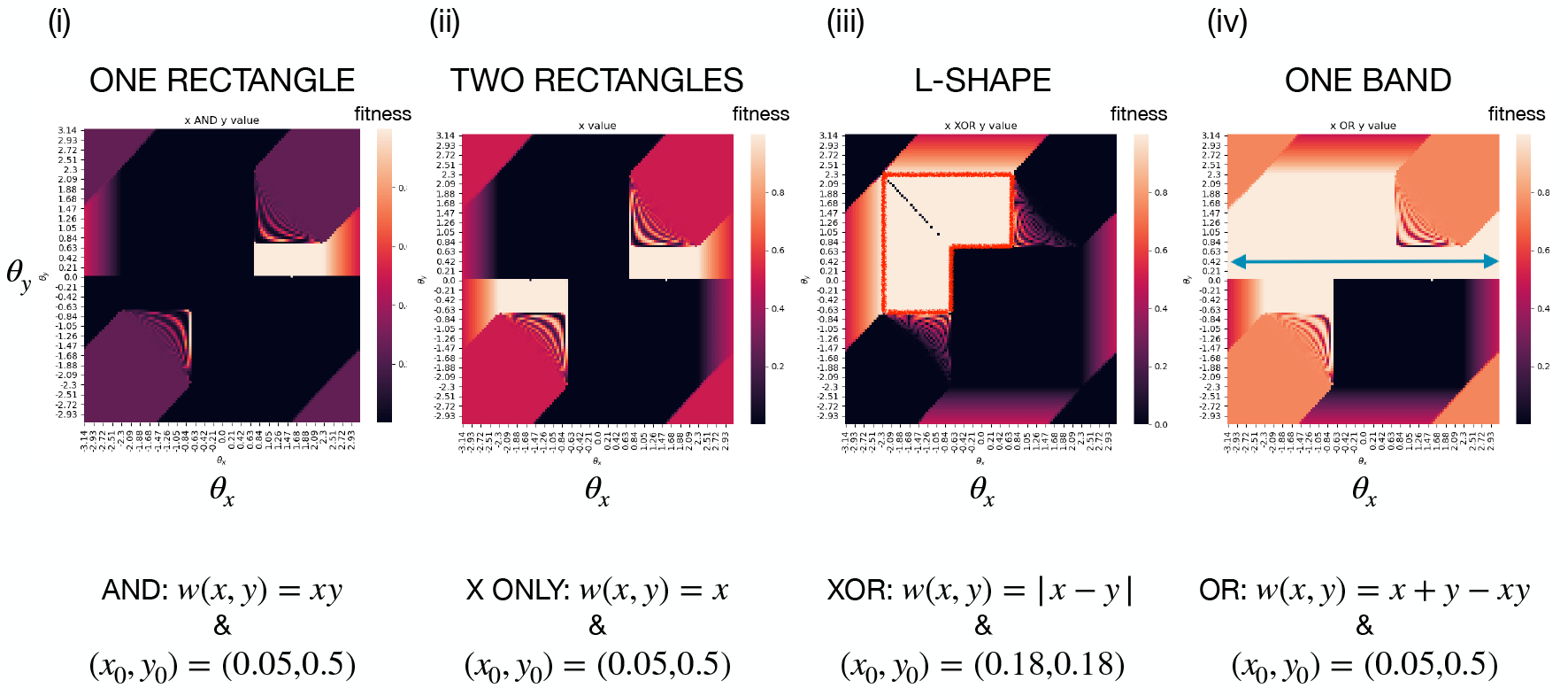
Four typical landscapes. The fitness value is represented by a color plotted as a function of (*θ*_*x*_, *θ*_*y*_). (i) A single rectangular region has maximum fitness. (ii) The maximum fitness region was divided into two parts. (iii) Even when the initial conditions were changed, the genotypes on the red line surrounding the region achieved maximum fitness. (iv) The arrowed band extends in the horizontal direction. If *θ*_*y*_ is fixed in this range, *θ*_*x*_ will be maximally fit for any value (neutral for *x*).

##### ONE RECTANGLE (seen in AND)

Fig.5(i) presents an example of the “rectangle” landscape, where the region of maximum fitness is in a single rectangle. This landscape is mainly observed in the AND fitness function. The high-fitness region occurs only in the parts of classes S and Q of the AND fitness because the optimal fitness was achieved for only (*x, y*) = (1, 1). In the Θ space for S in Fig.5(i) the maximum fitness is determined by the initial expression (*x*_0_, *y*_0_).

##### TWO RECTANGLES (seen in X ONLY)

This fitness landscape was composed of two rectangular high-fitness regions (Fig.5(ii)) where *x* = 1 and *y* values were *y* = 1 for one region and *y* = 0 for the other. The fitness maximum was X ONLY when a stable fixed point satisfied *x* = 1. The dynamics that satisfy this condition were either S or D and the Θ space for S and A in Fig.5(i) reached the maximum fitness, which was determined by the initial expression (*x*_0_, *y*_0_) = (0.05, 0.5). Q achieved maximum fitness, but all maximum fitness regions were connected.

##### L-SHAPE (seen in XOR or OR)

The region of maximum fitness was distributed in an L-shape, as shown in Fig.5(iii) with the XOR fitness function. The shape of this landscape was an inverted L, which we collectively referred to as an L shape. This landscape was obtained when the fitness function was either an XOR or an OR. When using the XOR or OR fitness functions, the initial condition (*x*_0_, *y*_0_) existed in 0 ≤ *x*_0_ ≤ 1 and 0 ≤ *y*_0_ ≤ 1, except for the unstable fixed-point singularity. We found that the region of L-SHAPE had the maximum fitness for all initial conditions tested. Therefore, the fitness was always maximum in the dynamics-type-A region because the stable fixed points of dynamics-type-A were only (1, 0) and (0, 1), respectively. The L-shaped area was robust against noise in the expression or initial conditions with XOR or OR fitness.

##### ONE BAND (seen in OR or X ONLY)

The maximum fitness region was extended to the entire range of *θ*_*x*_ or *θ*_*y*_ as a band for the OR fitness function, as shown in Fig.5(iv). This band extended over the boundary of the Θ torus in a single direction and therefore, the expression level of one *x, y* was neutral in this band. In OR, fitness was maximized where *x* = 1 or *y* = 1, which was when a stable fixed point was reached and the dynamic types S, D, Q, Cy, and Cx were satisfied. In particular, in dynamics-type-D, the fitness of the OR was always maximally independent of the initial condition because only the stable fixed points were (1,0) and (0,1), and there were no other steady states. In S, Q, Cy, and Cx, the maximum fitness regions were determined by the initial expression (*x*_0_, *y*_0_). Depending on the initial expression, this type of landscape was also observed in the X ONLY fitness landscape.

#### Genotype distribution is restricted by evolution with sexual reproduction

Thus far, we have investigated the shape and features of the fitness landscape by comprehensively calculating the correlations between genotype, phenotype, and fitness in a two-gene network. By introducing Θ as a genotype parameter, we were able to examine the phenotype and fitness for a given genotype and obtain the global structure of the fitness landscape. Once we have information on the global fitness landscape, we can predict the gene distribution and evolvability of the population during sexual reproduction and mutation. Thus, we investigated how the population distribution of the gene parameter Θ changed through evolution, with or without sexual reproduction.

In particular, we focused on how the population distribution was restricted to a convex set of maximum fitness regions by evolution during sexual reproduction (i.e., convex regionalization). We introduced asexual and sexual reproduction into this model and numerically evolved the population distribution in these conditions for each typical fitness landscape.

#### Definition of mutation and recombination

Before discussing convex regionalization, we defined the procedure for genetic evolution. We considered two inheritance modes: asexual and sexual reproduction processes. In asexual reproduction, GRN *J*^new^ in the next generation was chosen from a set of *J*^fit^ with high fitness in the population. For mutations, the network adjacency matrix for the next generation, *J*^new^, was changed by adding a random value generated by a normal distribution with mean 0 and variance *σ*. Note that the mutation was not introduced to Θ but to *J*^new^. This was because Θ was an abstract characteristic value and not the actual value for simplicity. In sexual reproduction, two individuals, *J*^fit1^ and *J*^fit2^, were selected from the set *J*^fit^ to generate high fitness. The GRN in the next generation was then defined by row-wise mixing of the two highly fitted individuals. Therefore,

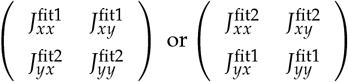

yields *J*^new^. This corresponded to the recombination of the promoter or enhancer regions. The evolutionary change in regulatory regions is much faster than in the gene that encodes for the protein itself (Luscombe *et al*. (2004)). Because proteins can bind to gene promotor regions, it is appropriate to use row-by-row mixing of the GRN adjacency matrix *J* as a model for sexual reproduction with free recombination. In the Θ torus, sexual reproduction could be expressed as parents with 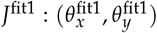 and 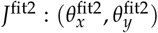, and with 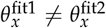 and 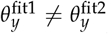. Additionally, suppose a rectangle with each side parallel to the *θ*_*x*_ or *θ*_*y*_ axis (Fig.6). By mixing the row vectors of the adjacency matrices with sexual reproduction, the children from 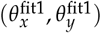 and 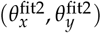 could be represented by 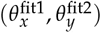 or 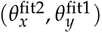. This corresponded to the vertex of the other side of the diagonal compared to the parent in the rectangle of Fig.6. Mutations in sexual reproduction were introduced in the same way as asexual reproduction, in which a normal distribution with mean 0 and variance *σ* were added to *J*^new^. Thus, sexual reproduction with a slight mutation could create a population of genotypes on the vertices of the rectangle.

**Figure 6.**
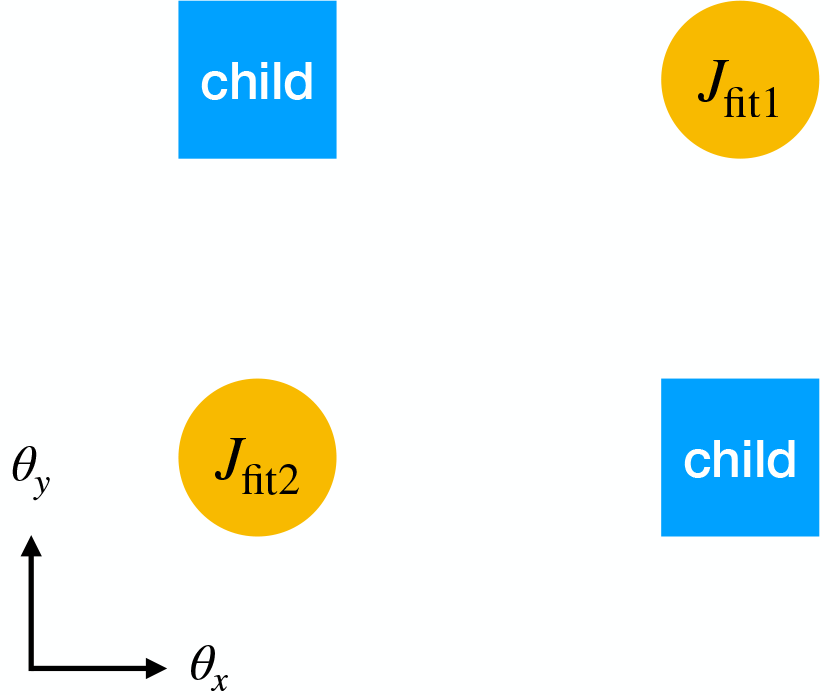
Transfer of genetic parameters (*θ*_*x*_, *θ*_*y*_) to children through recombination. Sexual reproduction involves mixing the row vectors of the adjacency matrix. The vector of children from the parents 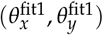 and 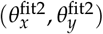 were randomly chosen from 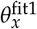 or 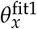, and 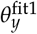 or 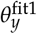. Graphically, two vertices can fall to a child on another diagonal from the parent rectangular axis.

#### Convex regionalization of sexually reproducing populations from a non-convex fitted area

For each fitness landscape category, we used simulations to examine changes in the population distribution due to sexual reproduction.

##### Simulation method

We ran simulations for 130 generations with a mutation size *σ* of 0.01 or 0.1, and a population size of 100. First, we set the initial expression to (*x*_0_, *y*_0_). The initial expression (*x*_0_, *y*_0_) was fixed and did not change throughout the subsequent simulations. Each element of *J*_*ij*_ in the 0th generation was given a uniform random number in the interval [-1,+1]. The dynamics of the GRN were calculated using Eq.(1) and Eq.(2) for 100 time steps. The average of each *x, y* in the last ten steps was used as the phenotype (expression). Fitness was calculated from the phenotype using the function defined in the previous section.

##### TWO RECTANGLES fitness landscape

The difference in the distribution between the asexually and sexually reproducing populations in the simulation is shown in Fig.7. During asexual reproduction, the population remained distributed in both maximum-fitness regions. In contrast, the evolution during sexual reproduction was concentrated only on one of the two fitted rectangles. These results can be explained as follows. The TWO RECTANGLES fitness landscape had two regions with equal maximum fitness values. If a population was distributed across. both, the offspring from the parent of the two rectangles fall onto the non-fitted region and the sexually reproducing population concentrated in one of the two regions was selected. Thus, the population distribution was in a convex region and sustained. In contrast, during asexual reproduction, the population was distributed across the two regions. Therefore, evolution of sexual reproduction could induce speciation. Here, the offspring from the two regions were less fit, resulting in hybrid sterility.

**Figure 7.**
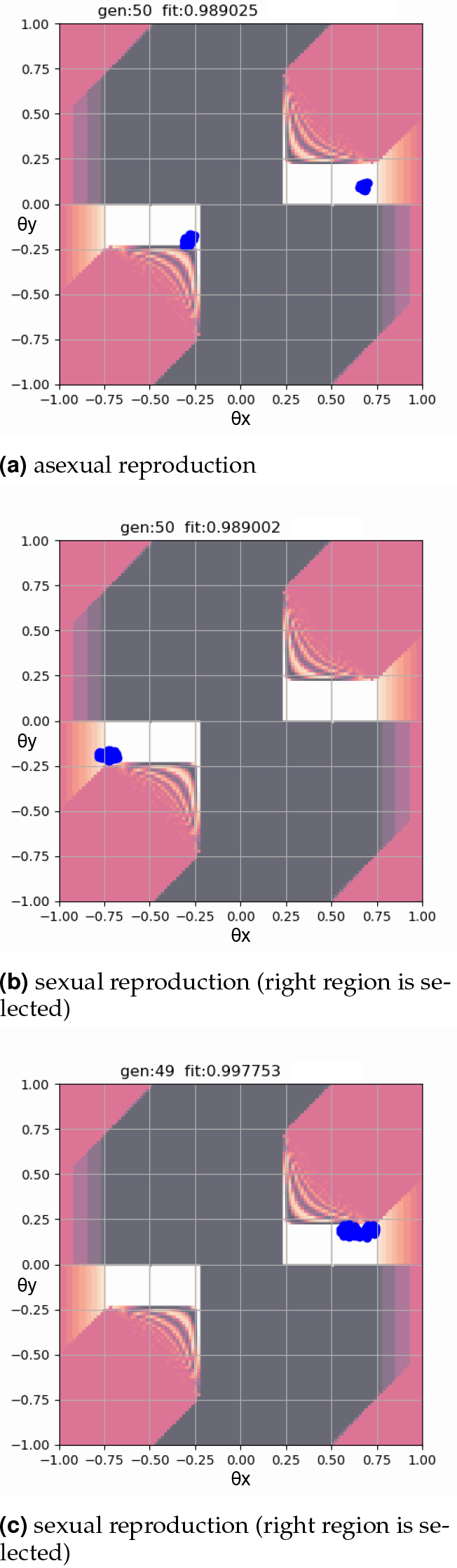
Comparison of the population distributions in two rectangular fitness landscapes with X ONLY(*w*(*x, y*) = *x*) fitness (a)during asexual reproduction (mutation only) or during sexual reproduction, where the population is branched into two cases, (b) and (c), which differ in each run of the evolution simulation. The initial expression was chosen as (*x*_0_, *y*_0_) = (0.05, 0.5). The mutation rate was set to 0.01. “gen:” number is the generation number and “fit:” number is the average fitness of the population.

Following the definition of speciation by hybrid sterility, it can be concluded that speciation occurred in this case.

##### L-SHAPE fitness landscape

The difference in population distribution between asexual and sexual reproduction was also observed in the L-SHAPE fitness landscape, as shown in Fig.8(iii). During asexual reproduction, the genotype population was spread over the entire L-shaped area of the landscape; however, during sexual reproduction, the population was biased in one direction of the L-SHAPE. Here, the L-SHAPE fitness landscape (Fig.9) extended vertically and horizontally in two directions, but the offspring from the parent between these two directions (IV of Fig.9) were less fit as a result of the genetic change in Fig.6.

**Figure 8.**
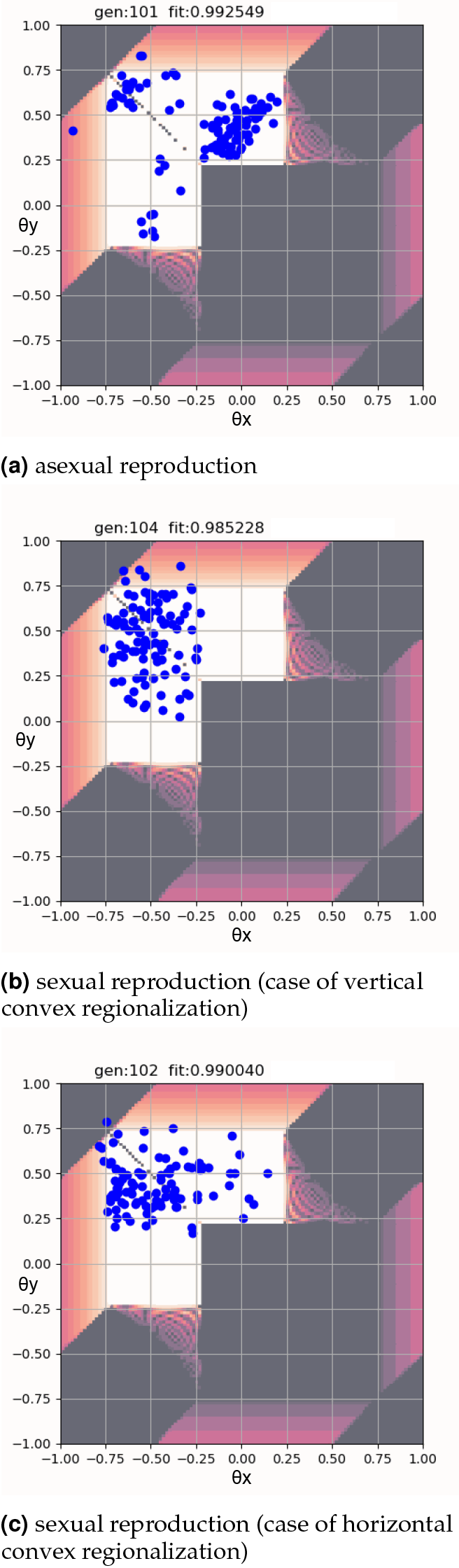
Comparison of the population distributions in two rectangular fitness landscapes with X ONLY(*w*(*x, y*) = *x*) fitness during (a) asexual reproduction (mutation only) and (b,c) sexual reproduction. The population branched into two cases, (b) and (c), which differ in each run of the evolution simulation. The initial expression was chosen as (*x*_0_, *y*_0_) = (0.18, 0.18). The mutation rate was set at 0.1. “gen:” number is the generation number and “fit:” number is the average fitness of the population.

**Figure 9.**
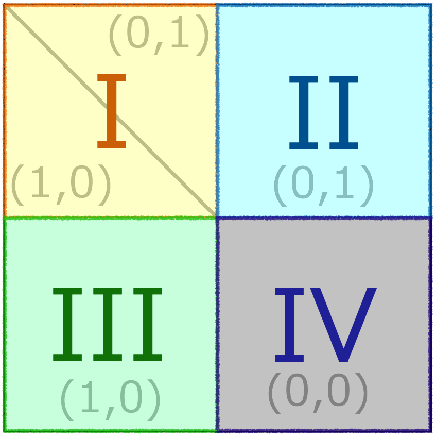
Partition of L-SHAPE fitness landscape. Regions I, II, and III support the highest fitness, while fitness is lost in region IV. Values with brackets like (1,0) shows the stable expression levels of *x* and *y*.

Hence, the offspring produced from sexual reproduction between the two branches of the L-SHAPE landscape (II and III in Fig.9) shrank into a rectangular region, either vertically (I and III in Fig.9) or horizontally (I and II in Fig.9). Such convex regionalization of the population distribution did not occur during asexual reproduction. This convex regionalization was similar to speciation in the Bateson-Dobzhansky-Muller model (Bateson (1909); Dobzhansky (1936, 1937); Muller (1940, 1942)), which is supposed to be an L-shaped fitness landscape. However, we found that some individuals from the common square area (I in Fig.9) of the two edges maintained high fitness and were not reproductively isolated, indicating that complete speciation did not occur. Only some of the two rectangular regions (II and III in Fig.9) were not fit, as an offspring could be located in IV in Fig.9. However, this convex regionalization was achieved so that the population in the L-SHAPE region was not allowed under sexual reproduction.

## Discussion

This study analyzed the genotype–phenotype relationship across all two-gene regulatory networks and obtained the fitness landscapes according to possible Boolean fitness functions. Characteristic landscapes including TWO RECTANGLES with the two optimal regions, L-SHAPE and ONE BAND, were obtained. As a result of evolution during sexual reproduction, the population becomes restricted to the convex region, leading in speciation.

To characterize the genetic changes in the gene regulatory matrix, a pair of continuous parameters Θ = (*θ*_*x*_, *θ*_*y*_) was introduced and the relationship between genotype and phenotype was characterized to examine how these relationships are associated with fitness and genome distribution. Several studies have discussed how the genotype-phenotype–fitness relationship affects GRN dynamics(Hether and Hohenlohe (2014); Friedlander *et al*. (2017); Schiffman and Ralph (2022)). However, these analyses were restricted to a specific level. For instance, genotype to phenotype in Hether and Hohenlohe (2014)), while a complete analysis of GRN dynamics has not been performed. We conducted a comprehensive analysis of the genotype–phenotype– fitness–distribution relationship. Furthermore, our results improve our understanding of how sexual reproduction changes population distribution or leads to speciation-like events in the fitness landscape, which requires a global landscape structure.

This study demonstrated that sexual reproduction limits populations only in a restricted, convex set in the maximum fitness regions (convex regionalization). The population was restricted to horizontal or vertical rectangular regions in the L-SHAPE landscape. In the TWO RECTANGLES landscape, the population was limited to one of the two separate high-fitness regions, which led to speciation. In the L-SHAPE landscape, complete speciation was not achieved, but convex regionalization led to two distinct population distributions, as in speciation.

When discussing the effects of multiple gene interactions, epistasis is often adopted. Epistasis is defined as a nonlinear change in fitness with multiple mutations. When the fitness change is lower or higher than the addition of changes in multiple mutations, it is called negative or positive epistasis, respectively. Epistasis is applied to local fitness changes, which benefits the study of the effects of relatively small mutations. In contrast, the fitness landscape provides global information on the fitness. This study showed that such information is essential for studying changes in population distribution, robustness of sexual reproduction, and speciation.

The method and results presented in this study can be used to solve other network related problems. First, the genotype parameter Θ may be used for evolution in other Boolean network systems such as machine learning and social or ecological networks(Raimundo *et al*. (2018); Saavedra *et al*. (2007); Shizuka and McDonald (2015); Sinha *et al*. (2022); Gordon (2014)). It can also be applied to a system that interacts with a steep sigmoid function. Here, the threshold *ζ* = 0.5 was used for simplicity and symmetry, but even if the threshold for each genes is changed, Θ can be used because the dynamics equivalent to this study were obtained by the transformation of variables, even though the classification of dynamics was more complex. In addition, Θ can be extended to systems with more than two (*N*) genes. In this case, the Θ space had *N*(*N* − 1) dimensions, which made it more difficult to obtain a global fitness landscape. However, the fixed points and their stability can be evaluated according to the value of Θ. The method of the present study can be applied by maintaining some Θ values within a certain range. For instance, the present results can be extended to a system of multiple pairs of corresponding genes. In general, by introducing network modules (network motifs Alon (2019)), the application of this method is straightforward.

In conclusion, our global analysis of GRNs based on Θ values and the characterized fitness landscape contributes to the comprehensive understanding of GRN evolution, particularly convex regionalization associated with sexual reproduction and resultant speciation.

## Acknowledgments

The authors would like to thank Tetsuhiro Hatakeyama, Yuichi Wakamoto, Naoki Irie, Akira Sasaki, and Shuji Ishihara for their stimulating discussions and Hideki Innan and Takahiro Sakamoto for providing information about genetics.

